# Analysis of multi-tissue transcriptomes reveals candidate genes and pathways influenced by cerebrovascular diseases

**DOI:** 10.1101/806893

**Authors:** Zhi-Lin Pan, Cho-Yi Chen

**Affiliations:** Institute of Biomedical Informatics, National Yang-Ming University, Taipei 112, Taiwan

## Abstract

Cerebrovascular diseases (CVD) are a group of medical conditions that impair circulation of blood to the brain, including stroke, transient ischemic attack (TIA), embolism, aneurysm, and other circulatory disorders affecting the brain. Here, we investigated the effects of having CVD history on the molecular signature of brain regions by comparing gene expression profiles from several brain tissues between cohorts with and without CVD history. We first merged tissue samples from GTEx RNA-Seq dataset into clusters based on the overall gene expression similarity. Then we performed differential expression (DE) analyses for each cluster using a linear mixed model that controls covariates and the individual random effect. Cross-region DE genes were ranked by the combined q-values derived from the mixed model using Fisher’s method. Functional enrichment analyses were performed using Gene Set Enrichment Analysis (GSEA) program. We identified hundreds of DE genes, and many of them are related to endothelial or brain functions and associated diseases. We found that STAB1 was highly overexpressed across brain regions in the CVD cohort, and the upregulation of STAB1 in brain tissues may contribute to weaker self-defense mechanisms against lesions in the brain. Our results suggest a list of candidate genes and pathways that may be dysregulated in the brains of people with CVD history, implying that suffering from CVD could pose potential hazard to the brain.

## Introduction

Cerebrovascular disease (CVD) remains one of the global leading causes of disability and mortality^1^, and the incidence and costs are projected to rise substantially in the future^2^. CVD is a group of clinical conditions that impair the blood flow to the brain, including stroke, transient ischemic attack (TIA), intracranial aneurysm, embolism, and other vessel diseases. It is a multifactorial and complex disease caused by genetic and environmental factors^1,3^. Previous genome-wide association studies (GWAS) and meta-analyses had identified various genetic loci associated with stroke in predominantly European-ancestry groups^4–25^, a recent multiancestry meta-analysis in 521,612 individuals discovered 22 new risk loci, bring up to 32. Most identified stroke loci are associated with other vascular traits and correlated with blood pressure^26^. Besides multiple genetic risk factors, several single-gene disorders also contribute to CVD^27–35^. However, genetic variability that contributes to susceptibility to these diseases and the mechanism underlying CVD largely remain to be identified.

Cerebrovascular diseases are also influenced by clinical features. Sex-related differences exist in incidence rates reported by community-based and hospital-based studies^36^. Age is another risk factor - the incidence and severity of stroke, the most significant entity of CVD, are significantly increased with age^37^. Likewise, the increase of body mass index (BMI) could lead to a significant increase in the relative risk of total stroke, ischemic and hemorrhagic stroke^38^.

Genotype-Tissue Expression (GTEx) project has established a database of clinical information such as medical histories, expression data, and whole-genome sequences, and helps discover the underlying genetic variability in multiple healthy human tissues^39^. The medical histories were from the hospital systems where recorded prior care of some deceased donors. Understanding normal physiology can provide new information on disease mechanisms and better treatment for patients.

In this study, we analyzed the 6,295 human transcriptomes covering 42 tissues from GTEx v6 release, to investigate the effects of having CVD medical history. We applied a linear model which corrects for confounding factors including sex, age, BMI, hardy scale, batch, and principal components inferred from genotypes to identify differentially expressed genes. Functional enrichment analyses reveal known biological functions and processes which are overrepresented in derived differential expressed genes. The main goal of stroke/CVD research is to develop effective treatments to reduce brain impairment from ischemic insult through a better understanding of the underlying pathogenic molecular mechanisms; although this study analyzed the expression data from normal tissues without true diseases, the results may let us know the long-term effects of CVD histories to the human body as well as deeper insights of CVD characteristics and mechanisms.

## Results

### Merging sample groups

The RNA-seq data from GTEx v6 release were downloaded from dbGaP. After preprocessing, normalizing and filtering the data as described in method, there are total 6295 samples (523 with and 5772 without CVD histories) covering 42 tissues from 428 subjects (41 with and 387 without CVD histories) (Supplementary Fig. S1).

In order to increase the statistical power by maximizing the sample sizes, we clustered transcriptionally indistinguishable tissue samples by performing principal component analysis (Supplementary Fig. S2). There are three main clusters of brain tissues – brain-0, brain-1 and brain-2. Since unequal subtissue composition in a cluster may lead to gene expression differences between the two cohorts, Chi-square tests were implemented to determine whether their composition is significantly different, and none of those P-values is greater than 0.05 (Table 1). Also, skin samples were grouped from lower leg (sun exposed) and from the suprapubic region (sun unexposed).

**Table 1.**
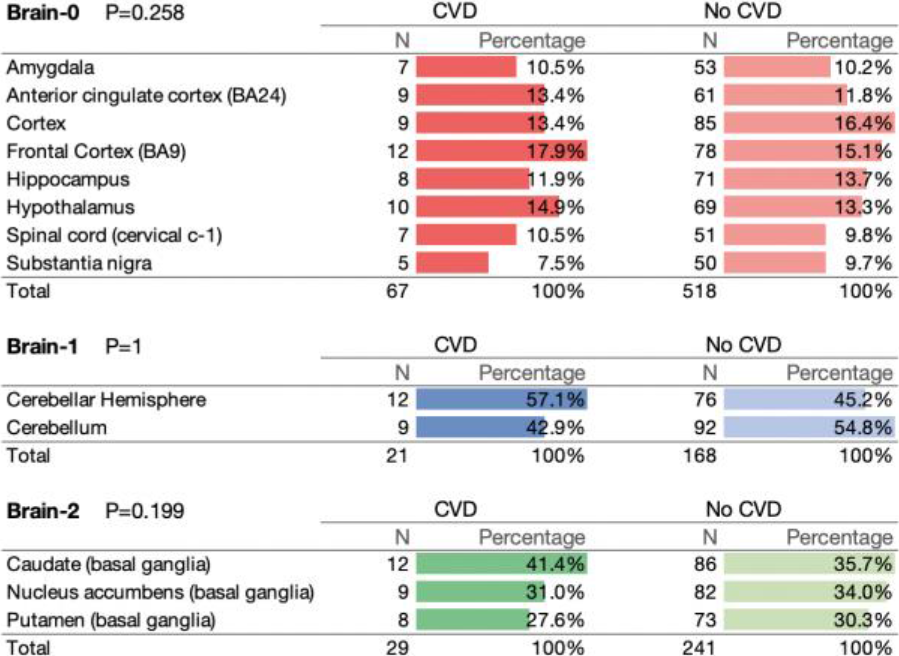
Tissue composition in brain clusters. P-values were calculated using chi-square test.

### Differential expression analysis of brain transcriptomes

Voom-limma^40,41^ pipeline was implemented to identify genes which are differentially expressed between cohorts with and without CVD histories. Details of the model see method. Genes at a false discovery rate (FDR) less than 0.1 and log2 fold change greater than 1.5 are considered significantly differential expressed (Supplementary Fig. S3).

For brain clusters, genes in limma reports were ranked by calculating Fisher’s combined P-value^42^ from the adjusted P-values in three clusters, showing a common set of genes which share similar expression patterns across brain regions. We yielded 220 significant combined differentially expressed (DE) genes (Fisher’s P-value < 0.1) and directions of t-statistics of 217 genes are same (Fig. 1). Among these genes with consistent directions, 125 are up-regulated while 92 are down-regulated (Supplementary Data S1).

**Figure 1.**
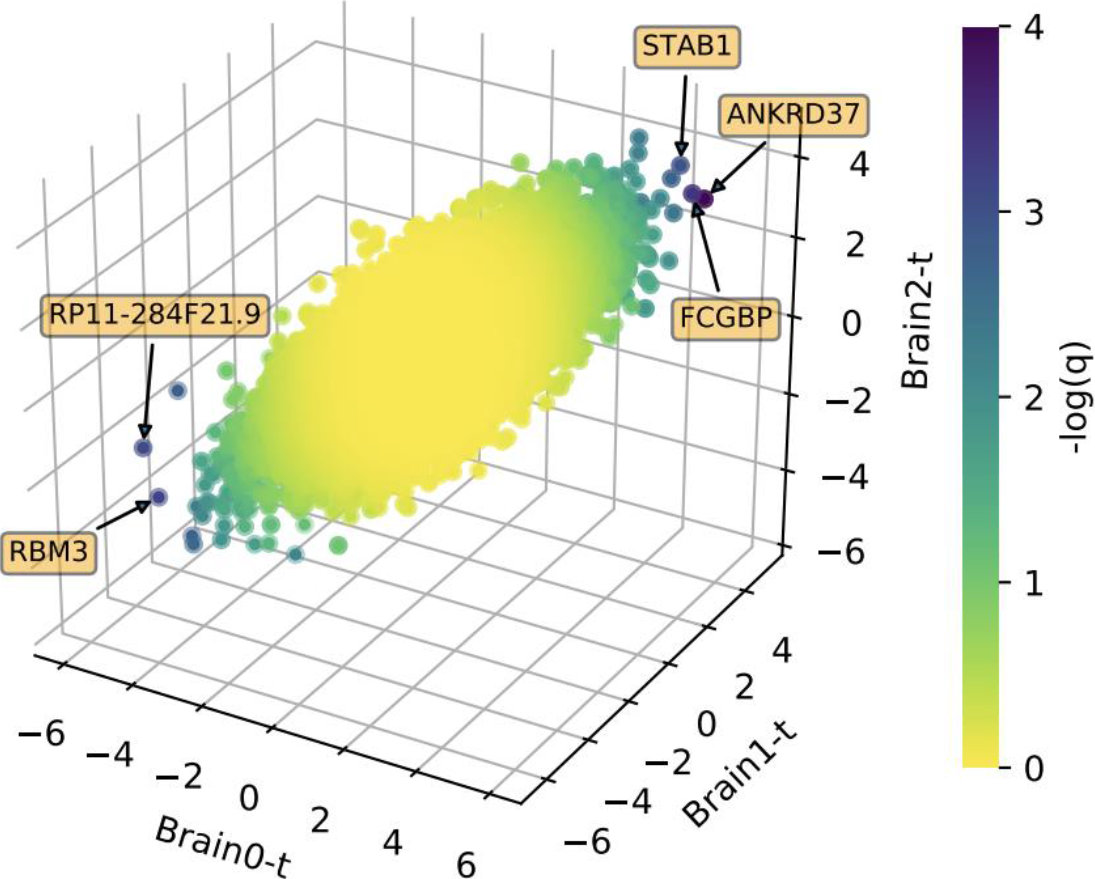
Scatter plot of t-statistics from three clusters. Each dot is expression value of one gene. Colors represent negative logarithm of combined Fisher’s q-values. Top 5 significant genes are highlighted within orange boxes.

Dozens of consistently significant DE genes are related to hypoxia response, and hypoxia refers to oxygen levels below normal and is a common occurrence following stroke. Some of these genes interact with hypoxia inducible factors-1 (HIF-1), one of the critical regulators of cellular response to hypoxia. For example, the most significantly upregulated gene *ANKRD37* (Fig. XA), *PFKFB3*, *FECH*, *GPRC5A*, *VEGFA*. Note that *VEGFA* (vascular endothelial growth factor A), which is critical for blood vessel growth in central nervous system (CNS) and is also triggered by hypoxia and inflammation, only significantly overexpressed in brain-0. Another top significant DE gene *RBM3* (Fig.XB), a RNA-binding protein induced by hypoxia and is neuroprotective under stress condition, however is downregulated in our DE results. None of the HIF genes were differentially expressed, maybe because the activation of HIF-1 only lasts for a period after cerebral ischemia^43^. A significant downregulated lncRNA in brain-0 and brain-1, *RP11-284F21.9* (Fig.XC), is antisense to *BCAN*^44^. BCAN (Brevican) encodes a member of proteoglycans that is specifically expressed in CNS, maintaining the extracellular environment of mature brain^45^. Among those upregulated genes, *SERPINA1* (Fig. XD), *TGFB1, TSPAN2* had been associated with subtypes of stroke by GWAS and meta-GWAS. In summary, our DE analysis on brain transcriptomes revealed genes related to occurrences after cerebrovascular accidents like the hypoxia mentioned above, including three genes previously implicated in stroke.

**Figure 2.**
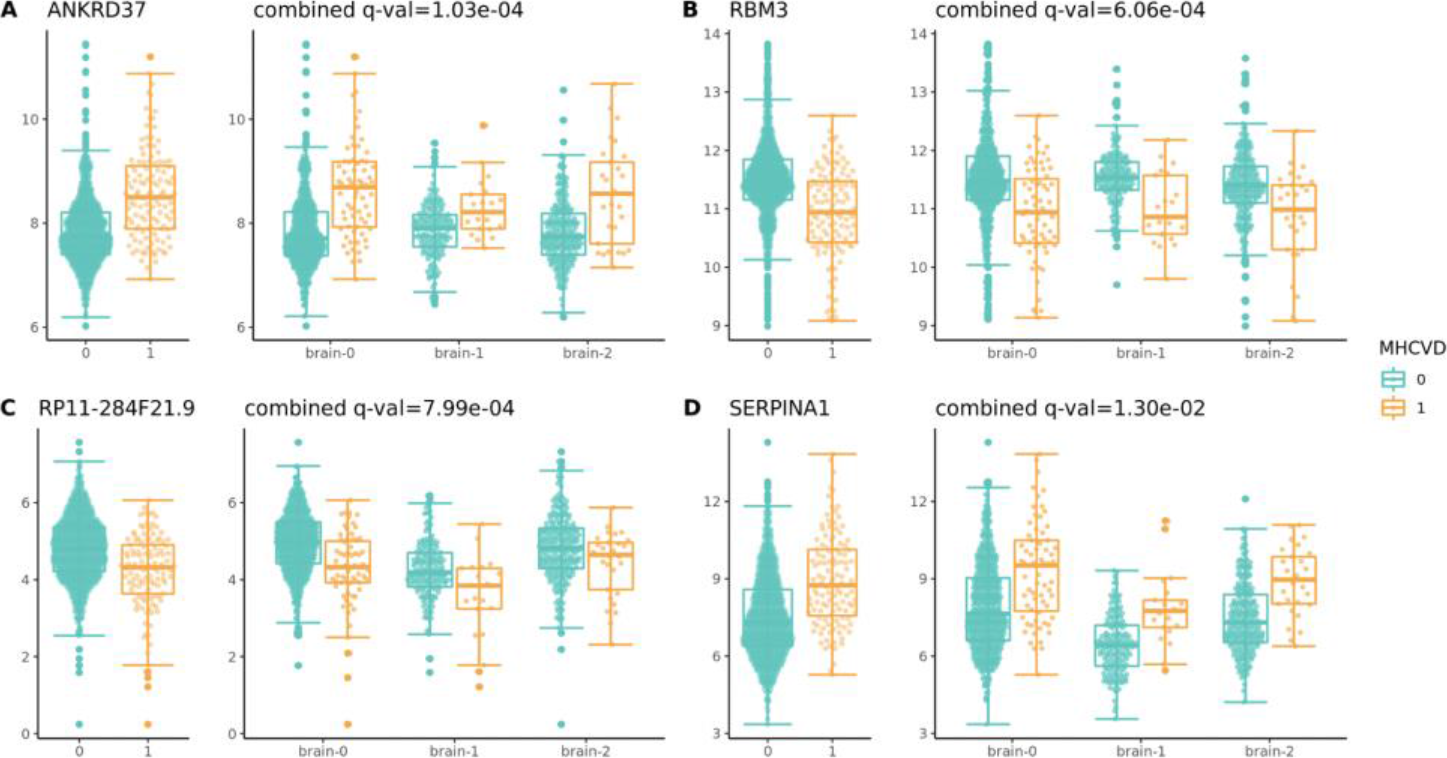
Top significant DE genes in brain groups. The left panel is the overall expression level in all groups, and the right panel is expression level in each group. Combined q-value is calculated using Fisher’s method.

**Table 2.**
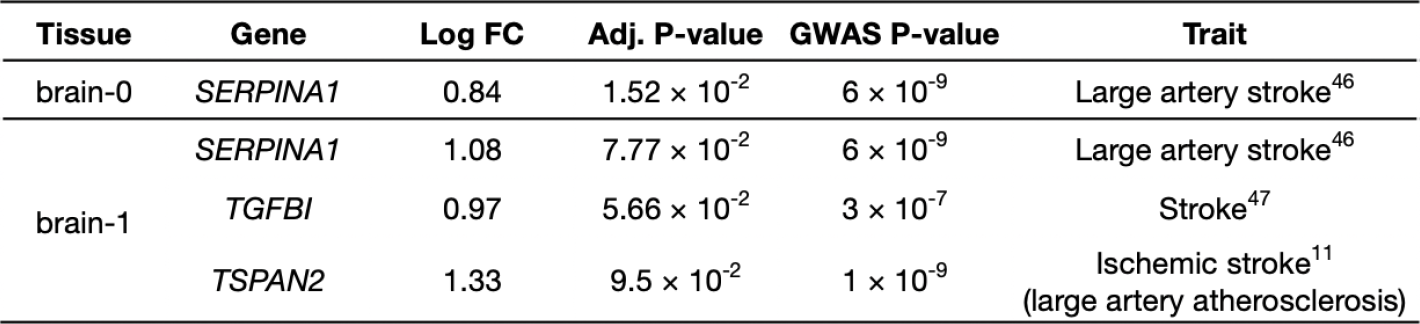
Summary of the DE genes which have been associated with stroke or subtypes of stroke by GWAS. The log2 fold change and adjusted p-value for each gene in the tissue, and the p-value from GWAS results were included.

DE genes were further analyzed by Gene Set Enrichment Analysis (GSEA) to gain a functional overview. GSEA presented a wide spectrum of Gene Ontology biological process (GOBP) terms, most top significantly enriched terms are shown in Fig 3. Generally, inflammatory mechanisms are significantly upregulated since most of the terms are linked to inflammation. Mast cells are perivascular resident cells distributed in most tissues around blood vessels and can also be found in central nervous system (CNS), they have been associated to various neuroinflammatory conditions of CNS like stroke^46^; the pro-inflammatory interferon gamma signaling is directly associated with stroke by neurodegeneration^47^; Toll-like receptors (TLRs) are master regulators of innate immunity and play an integral role in the activation of the inflammatory response during infections, also play a modulating role in ischemic brain damage after stroke^48^; proinflammatory cytokines, such as IL-6, have been implicated in several mechanisms that might promote ischemic brain injury^49^; and immune cell proliferation, differentiation, activation and cell adhesion are all involved in innate and adapt immune responses, then these responses as well as angiogenesis and coagulation share extensive cross talk with inflammatory reactions. Additionally, reactive oxygen species (ROS) metabolic process and apoptotic process are upregulated, not shown in the figure though. On the other hand, downregulated genes are enriched in several metabolic processes, mitochondrial functions and cell cycle events. Furthermore, “refined” hallmark gene sets enrichment reveals more specific pathways associated with neuroprotective effects (Supplementary Fig. S4), such as IL-6/JAK/STAT3, PI3K/AKT/mTOR, mTORC1 signaling.

**Figure 3.**
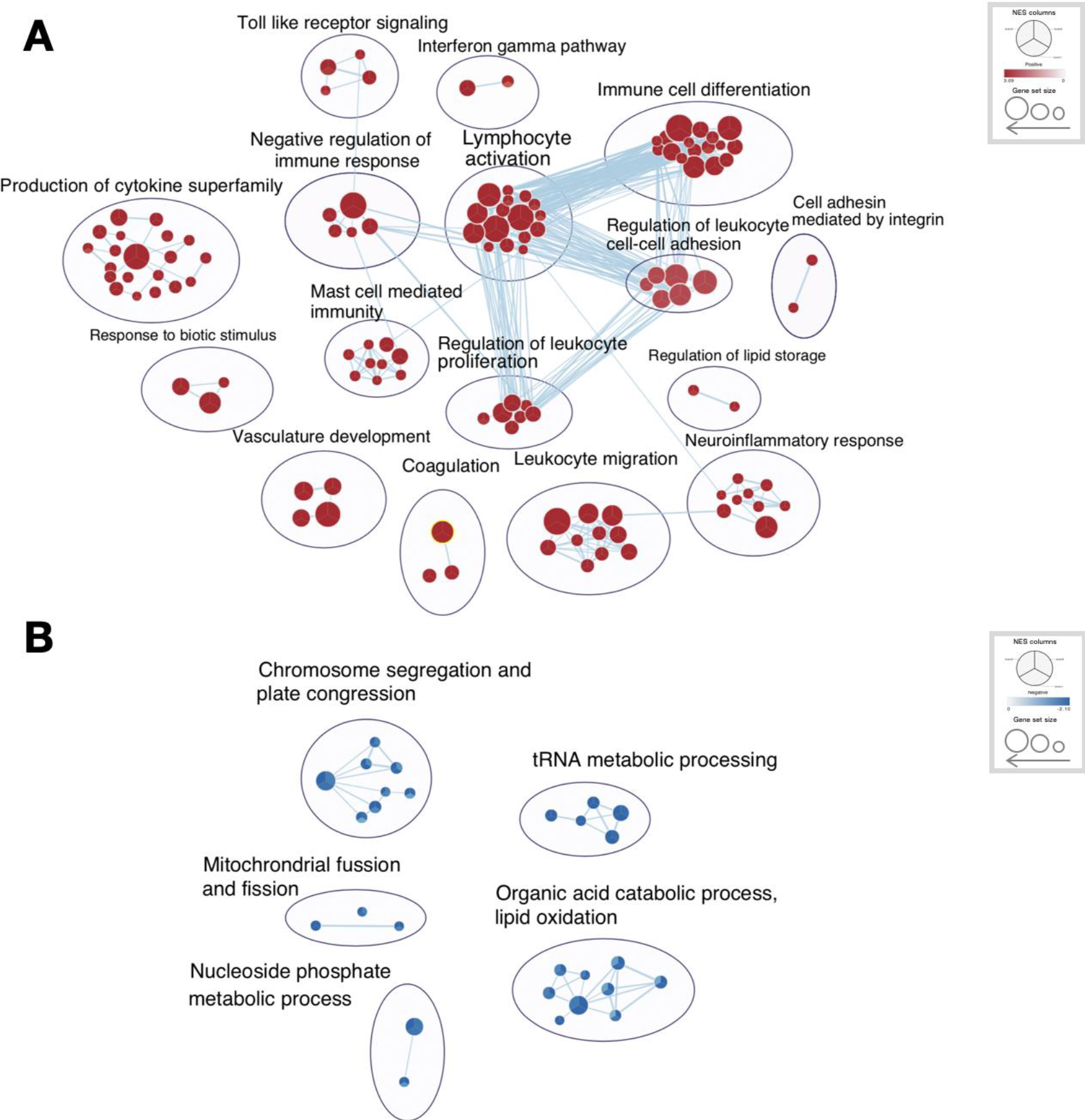
Enrichment map shows significantly enriched GOBP terms for genes upregulated and downregulated in the cohort with CVD history. Each circle composed with three parts which represent the P-values for each term in three brain clusters, the size of a circle correlated to the NES scroe.

Unsurprisingly, disease and phenotype associations are enriched in typical CVD-related traits: transient cerebral ischemia, intracranial aneurysm, atrial fibrillation, etc. and some vascular diseases like Ehlers-Danlos syndrome, Behcet’s disease and vasculitis (Supplementary Data X). Likewise, many autoimmune diseases, such as rheumatoid arthritis, systemic lupus erythematosus and immune thrombocytopenic purpura, as well as some other common diseases like Crohn’s disease and chronic obstructive pulmonary disease (COPD) are mapped to upregulated terms. These diseases may share common risk factors with cerebrovascular diseases. What’s more, infections or infectious diseases terms are also highly enriched, maybe due to infections are one of the common medical complication after stroke, leading to unfavorable functional outcome. Overall, these functional enrichment results link to the clinical conditions of CVD, common occurrences and complication after stroke as well as diseases not just in brain, and provide the possibly molecular evidence.

### Differential expression analysis of other tissues

On top of brain, several tissues - tibial artery, atrial appendage, minor salivary gland and skeletal muscle, skin - are also identified dozens or hundreds of DE genes by our linear model. Surprisingly, atrial appendage, actually the right atrial appendage (RAA) collected by GTEx (SMSMPSTE, phv00169238.v6.p1.c1), has over 1,000 significant DEGs. Interestingly, there is no information on relationship between RAA and stroke, only a weak incidence of atrial thrombosis^50^, while left atrial appendage (LAA) is the site most commonly correlated with thrombus formation that might increase risk of stroke^66^.

Multiple genes identified by stroke and aneurysm GWAS are overlapped with DE genes in some tissues (Table 3). For instance, *ALDH2* which associates with ischemic stroke and intracranial aneurysm is downregulated in visceral adipose (Fig. 4). Genetic variants in *ALDH2* exhibit association with visceral fat accumulation in male^51^ and hypertension^52^, both mediated by alcohol consumption, and lead to lower gene expression levels in adipose tissue. Increased visceral adipose tissue (VAT) is a risk factor for carotid artery atherosclerosis, atrial fibrillation and stroke, and low VAT proportion associates with better outcomes in acute ischemic stroke patients^53^.

**Table 3.**
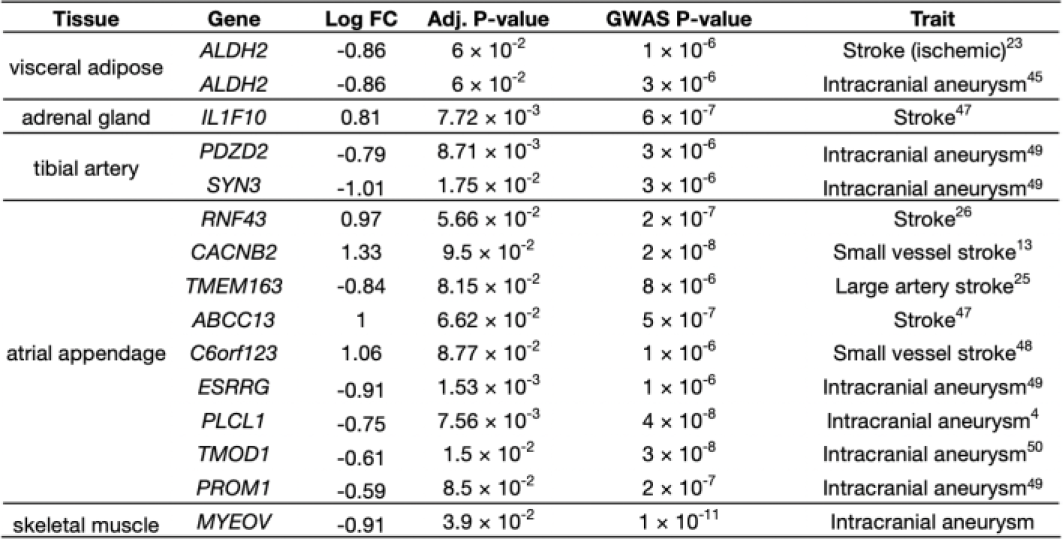
Summary of the DE genes which have been associated with stroke and aneurysm by GWAS. The log2 fold change and adjusted p-value for each gene in the tissue, and the p-value from GWAS results were included.

**Figure 4.**
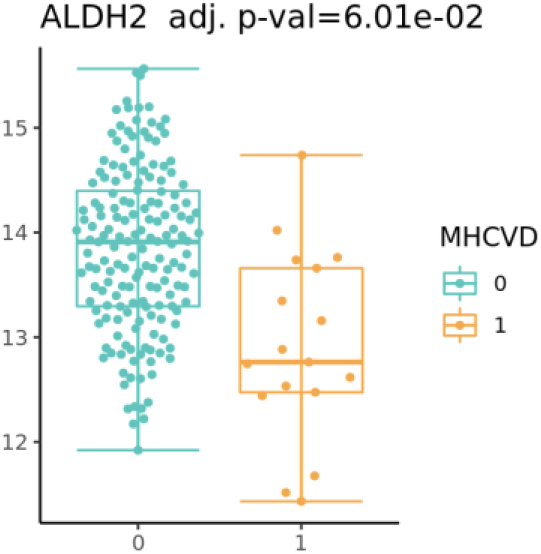
Expression level of DE gene overlapped with GWAS results *ALDH2* in visceral adipose

Almost all these tissues are enriched in mitochondrial-related terms, for instance, mitochondrion organization, mitochondrial electron transport and cellular respiration (Supp Table X). These terms are downregulated in visceral adipose, atrial appendage, skeletal muscle, pituitary and skin, while upregulated in tibial artery and minor salivary gland. Concomitantly, metabolic and catabolic processes are also downregulated in some tissues but upregulated in other tissues. Moreover, enriched muscle or cardiac muscle contraction terms are mapped to atrial appendage, tibial artery and skeletal muscle. In summary, ——.

## Discussion

Large-scale genome-wide association studies have obtained more and more genetic loci related to cerebrovascular disease with the increased amounts of genetic data and curated databases. However, molecular changes on mRNA level is difficult to prove and most transcriptome analyses on CVD only restricted to middle cerebral artery occlusion (MCAO) mouse and rat model or human blood expression data due to the limitation of tissue accession. Here, we analyzed multiple human tissue expression data from GTEx, although the condition is just having or not having CVD medical history also samples were all reviewed as normal, dozens to thousands of DE genes were identified across some of the tissues. Therefore, our results may indicate and prove what complications and events would occur after CVD and even what contributes to CVD causes, revealing underlying molecular signatures and mechanisms and providing new perspectives for further CVD studies.

We first built a linear mixed model, which allows both fixed and random effects, and applied it to expression data. The results of functional enrichment analysis based on identified differentially expressed genes successfully present most of the conditions after cerebrovascular diseases. Take ischemic stroke as an example, briefly, brain ischemia triggers inflammation followed by the generation of reactive oxygen species (ROS), these initiators of inflammation will activate microglia, the resident immune cell in brain. Next, microglia produce more pro-inflammatory cytokines leading to adhesion molecules induction in the cerebral vasculature. Cytokines then infiltrate more immune cells into ischemic brain, and inflammatory cells release cytotoxic agents such as nitric oxide (NO) causing brain cell damage exacerbation as well as the disruption of the extracellular matrix and blood-brain barrier (BBB)^54,55^. Therefore, therapies targeting both acute and chronic neuroinflammation could effectively reduce further brain damage and lessen long-term effects on neurological function.

Hypoxia is also closely associated with inflammatory condition and it causes cell injury and induce microglial activation, since many genes regulated by hypoxia involved in inflammatory responses^56^. Hypoxia following stroke is common and often attributed to pneumonia, aspiration and respiratory muscle dysfunction, with some other related syndromes^57^. We found that *ANKRD37* (Ankyrin Repeat Domain 37) was highly overexpressed across brain regions in cohort with CVD history. *ANKRD37* is induced in hypoxia and it is targeted by transcription factor Hypoxia-inducible factor 1 (HIF-1)^58^. It may be a clue about how hypoxia involves in brain damage following CVD, while *ANKRD37* is weakly documented, further experiments and analysis are necessary. Similarly, several genes are worth being notice, such as *RBM3* and *RP11-284F21.9* mentioned above and another upregulated gene *STAB1*. *STAB1* encodes a transmembrane receptor protein that may function in angiogenesis, lymphocyte homing, cell adhesion, defense against bacterial infection, or receptor scavenging for acetylated low density lipoprotein.

Moreover, genetic epidemiology reveals genetic variants associated with stroke. *ALDH2*-rs671, the variant associated with alcohol intake, has been used to conduct stroke mendelian randomization studies, proving alcohol consumption uniformly increases blood pressure and stroke risk^59^. And in our results, *ALDH2* indeed downregulated in visceral adipose tissue which is well in line with previous reports. However, the associations were only replicated in male Asian population, there may be more details of mechanisms of *ALDH2* to be explored. Our results may provide clues for further genetic studies. When we investigated the DE genes across analyzed tissues, we found excessive numbers of genes identified in right atrial appendage (RAA), the entire anterolateral triangular part of the right atrium. Despite most interests of stroke prevention (i.e. left atrial appendage closure) in atrial fibrillation (AF) patients are on left atrial appendage (LAA), the role of RAA in disease risk need to be elucidate and RAA might be an alternative position for stroke or cardiovascular diseases prevention.

The DTHFUCOD First Underlying Cause Of Death may affect the results

## Materials & Methods

### GTEx data

We collected RNA-Seq data from the Genotype-Tissue Expression (GTEx) project v6 release^39^ (dbGaP: phs000424.v6.p1). The RNA-Seq data were filtered and preprocessed using Yet Another RNA Normalization pipeline (YARN) R package^67^, and normalized in a tissue-aware manner by smooth quantile normalization^68^. GTEX-11ILO was remove due to potential sex misannotation. We only selected subjects that had MHCVD value 0 or 1 and genotype PCs, also excluded sex-specific tissues (ovary, prostate, testis, uterus, vagina) and cell lines. Moreover, the suboptimal samples, those with “FLAGGED” in the SMTORMVE field, were removed. Finally, total 6,295 samples were included in downstream analysis. Then, based on the similarity of expression profiles, we grouped 13 brain subregions into three main clusters by PCA: Brain-0 (amygdala, anterior cingulate cortex, frontal cortex, hippocampus, hypothalamus, spinal cord, substantia nigra), Brain-1 (cerebellar hemisphere, cerebellum) and Brain-2 (caudate, nucleus accumbens, putamen).

The medical history of cerebrovascular disease of GTEx subjects was recorded as value 0 (No), 1 (Yes) and 99 (Unknown) in variable MHCVD (dbGaP: phv00169142.v6.p1). The first 20 genotype principal components (PCs) of 450 donors were downloaded from dbGaP (phs000424.v6.p1), and the top three PCs were used as they captured the major genotype variance.

### Differential expression analysis

Differential expression (DE) analysis between CVD and non-CVD samples was conducted using the voom-limma pipeline^40,41^: RNA-Seq read counts were transformed to log counts per million (log-cpm) with associated precision weights to stabilize the variance in the data using the voom function, followed by linear model fitting and empirical Bayes procedure. In each brain tissue cluster, we modelled gene expression using the following linear regression model with blocking due to multiple and various tissue sites:

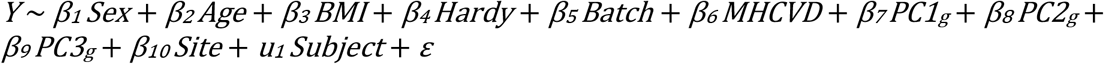

where *Y* is the gene expression level; *Sex* denotes the reported sex of the subject; *Age* denotes the age of the subject; *BMI* denotes the body mass index of the subject; *Hardy* denotes the 4-point Hardy scale of the subject (death classification), subjects where Hardy scale was missing were excluded; *Batch* denotes the type of nucleic acid isolation batch of the sample; *MHCVD* denotes whether the subject had CVD medical history; *PCg1~3* denote the first three PC values inferred from the subject genotype; *Site* denotes the tissue type of the sample; *Subject* denotes subject ID of the sample and was treated as a random effect factor.

In each tissue except brain tissues, we adopted the following linear regression model without blocking:

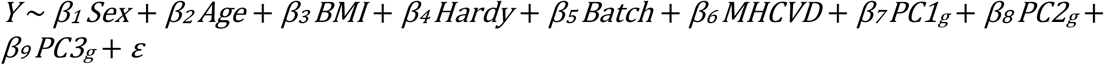

To control the false discovery rate (FDR), the P-values from the regression model were adjusted for multiple testing using Benjamini-Hochberg method^69^. The DE genes of brain clusters were ranked by the combined adjusted P-values derived from the three adjusted P-values in the results of each cluster using Fisher’s method^42^. The significant threshold of DE genes was set as 0.1 FDR.

### Functional Enrichment Analysis

Pre-ranked Gene Set Enrichment Analysis (GSEA)^70^ was conducted with gene lists ranked by the *t*-statistics from the results of DE analysis with default program parameters and a default background set on GSEA v4.0.1. Gene Matrix Transposed (GMT) files of Gene Ontology (GO) were downloaded from Molecular Signatures Database v7.0. Disease Ontology (DO) annotation file was downloaded from Alliance of Genome Resource and processed to gmt file. Likewise, Human Phenotype Ontology (HPO) gmt were modified from the file downloaded on HPO website^71^.

### Collection of costumed genes

GWAS association data were all downloaded from GWAS catalog using keyword “stroke” and “brain aneurysm”.

---

**Supplementary Fig. S1.**
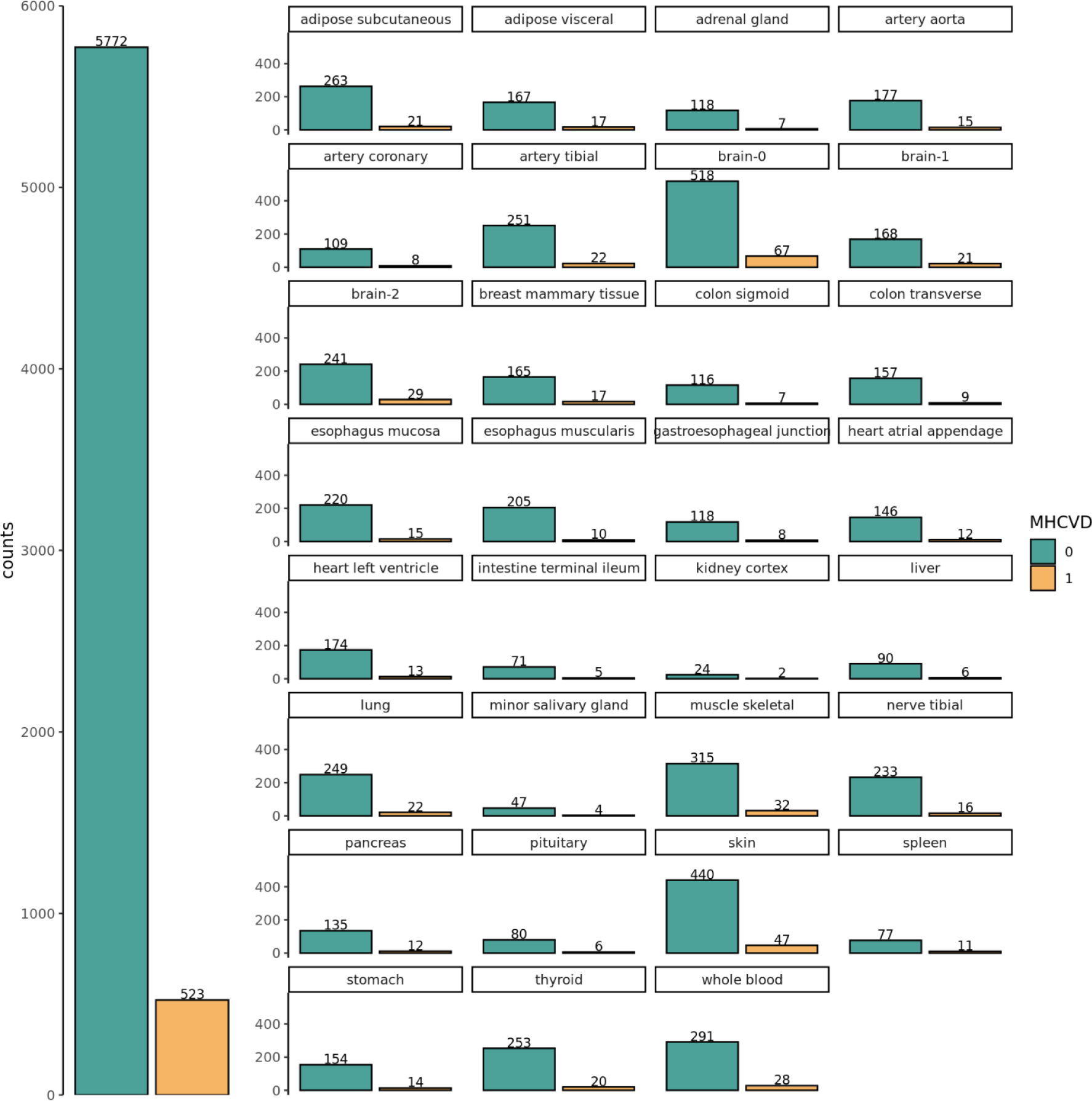
Number of samples with and without CVD history in each selected tissue and cluster.

**Supplementary Figure S2.**
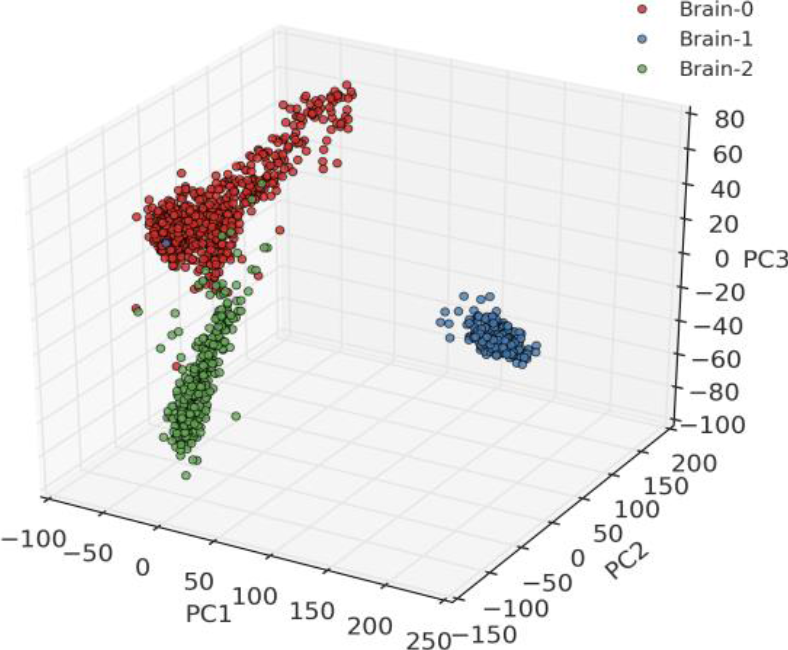
PCA plot shows three clusters among the samples of brain tissues in the gene expression space.

**Supplementary Figure S3.**
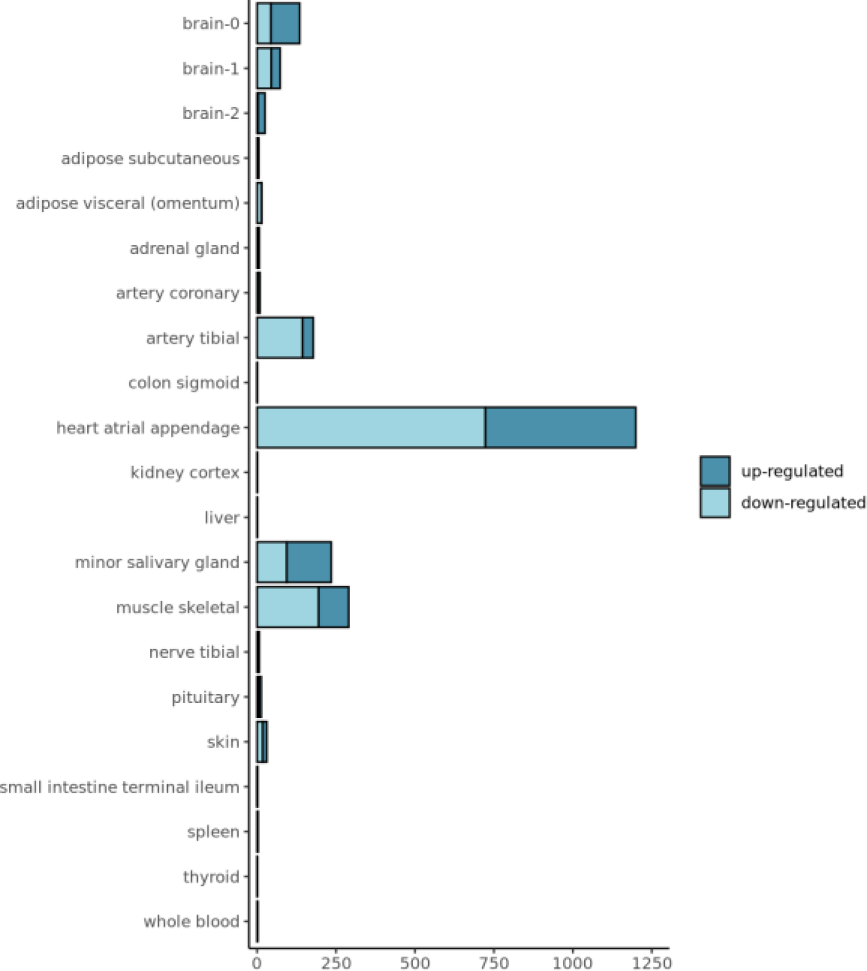
Number of significant genes identified by our linear regression model in each tissue and cluster. Significant means FDR < 0.1 and log2 fold change > 1.5.

**Supplementary Figure S4.**
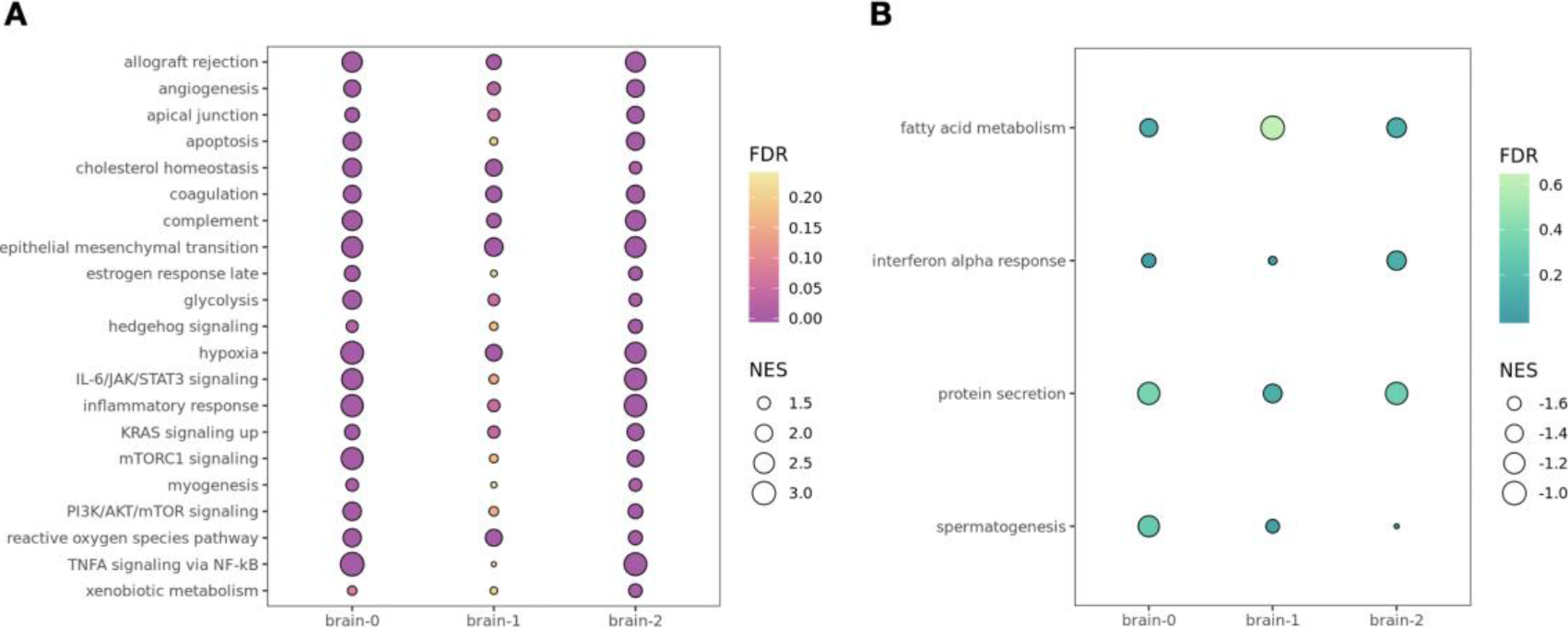
GSEA results for hallmark gene sets. The color and the size correlate to the FDR values and NES score.

## References

1. Benjamin, E. J. et al. Heart Disease and Stroke Statistics-2018 Update: A Report From the American Heart Association. Circulation (2018). doi:10.1161/CIR.0000000000000558

2. Ovbiagele, B. et al. Forecasting the future of stroke in the united states: A policy statement from the American heart association and American stroke association. Stroke (2013). doi:10.1161/STR.0b013e31829734f2

3. Boehme, A. K., Esenwa, C. & Elkind, M. S. V. Stroke Risk Factors, Genetics, and Prevention. Circulation Research (2017). doi:10.1161/CIRCRESAHA.116.308398

4. Bilguvar, K. et al. Susceptibility loci for intracranial aneurysm in European and Japanese populations. Nat. Genet. (2008). doi:10.1038/ng.240

5. Ikram, M. A. et al. Genomewide association studies of stroke. N. Engl. J. Med. (2009). doi:10.1056/NEJMoa0900094

6. Woo, D. et al. Meta-analysis of genome-wide association studies identifies 1q22 as a susceptibility locus for intracerebral hemorrhage. Am. J. Hum. Genet. (2014). doi:10.1016/j.ajhg.2014.02.012

7. Williams, S. R. et al. Genome-Wide Meta-Analysis of Homocysteine and Methionine Metabolism Identifies Five One Carbon Metabolism Loci and a Novel Association of ALDH1L1 with Ischemic Stroke. PLoS Genet. (2014). doi:10.1371/journal.pgen.1004214

8. Traylor, M. et al. A Novel MMP12 Locus Is Associated with Large Artery Atherosclerotic Stroke Using a Genome-Wide Age-at-Onset Informed Approach. PLoS Genet. (2014). doi:10.1371/journal.pgen.1004469

9. Foroud, T. et al. Genome-wide association study of intracranial aneurysm identifies a new association on chromosome 7. Stroke (2014). doi:10.1161/STROKEAHA.114.006096

10. Rannikmäe, K. et al. Common variation in COL4A1/COL4A2 is associated with sporadic cerebral small vessel disease. Neurology (2015). doi:10.1212/WNL.0000000000001309

11. NINDS Stroke Genetics Network (SiGN), I. S. G. C. (ISGC). Loci associated with ischaemic stroke and its subtypes (SiGN): a genome-wide association study. Lancet. Neurol. (2016). doi:10.1016/S1474-4422(15)00338-5

12. Williams, S. R. et al. Shared genetic susceptibility of vascular-related biomarkers with ischemic and recurrent stroke. Neurology (2016). doi:10.1212/WNL.0000000000002319

13. Cheng, Y. C. et al. Genome-Wide Association Analysis of Young-Onset Stroke Identifies a Locus on Chromosome 10q25 Near HABP2. Stroke (2016). doi:10.1161/STROKEAHA.115.011328

14. Hinds, D. A. et al. Genome-wide association analysis of self-reported events in 6135 individuals and 252 827 controls identifies 8 loci associated with thrombosis. Hum. Mol. Genet. (2016). doi:10.1093/hmg/ddw037

15. Traylor, M. et al. Shared genetic contribution to ischemic stroke and Alzheimer’s disease. Ann. Neurol. (2016). doi:10.1002/ana.24621

16. Gudbjartsson, D. F. et al. A sequence variant in ZFHX3 on 16q22 associates with a trial fibrillation and ischemic stroke. Nat. Genet. (2009). doi:10.1038/ng.417

17. Working, N. et al. Identification of additional risk loci for stroke and small vessel disease: a meta-analysis of genome-wide association studies. Lancet. Neurol. (2016). doi:10.1016/S1474-4422(16)00102-2

18. Traylor, M. et al. Genetic variation at 16q24.2 is associated with small vessel stroke. Ann. Neurol. (2017). doi:10.1002/ana.24840

19. Yasuno, K. et al. Genome-wide association study of intracranial aneurysm identifies three new risk loci. Nat. Genet. (2010). doi:10.1038/ng.563

20. Bellenguez, C. et al. Genome-wide association study identifies a variant in HDAC9 associated with large vessel ischemic stroke. Nat. Genet. (2012). doi:10.1038/ng.1081

21. Holliday, E. G. et al. Common variants at 6p21.1 are associated with large artery atherosclerotic stroke. Nat. Genet. (2012). doi:10.1038/ng.2397

22. Foroud, T. et al. Genome-wide association study of intracranial aneurysms confirms role of anril and SOX17 in disease risk. Stroke (2012). doi:10.1161/STROKEAHA.112.656397

23. Traylor, M. et al. Genetic risk factors for ischaemic stroke and its subtypes (the METASTROKE Collaboration): A meta-analysis of genome-wide association studies. Lancet Neurol. (2012). doi:10.1016/S1474-4422(12)70234-X

24. Williams, F. M. K. et al. Ischemic stroke is associated with the ABO locus: the EuroCLOT study. Ann. Neurol. (2013). doi:10.1002/ana.23838

25. Dichgans, M. et al. Shared genetic susceptibility to ischemic stroke and coronary artery disease: a genome-wide analysis of common variants. Stroke. (2014). doi:10.1161/STROKEAHA.113.002707

26. Malik, R. et al. Multiancestry genome-wide association study of 520,000 subjects identifies 32 loci associated with stroke and stroke subtypes. Nat. Genet. (2018). doi:10.1038/s41588-018-0058-3

27. Joutel, A. et al. The ectodomain of the Notch3 receptor accumulates within the cerebrovasculature of CADASIL patients. J. Clin. Invest. (2000). doi:10.1172/JCI8047

28. Hara, K. et al. Association of HTRA1 mutations and familial ischemic cerebral small-vessel disease. N. Engl. J. Med. (2009). doi:10.1056/NEJMoa0801560

29. Grabowski, T. J., Cho, H. S., Vonsattel, J. P. G., William Rebeck, G. & Greenberg, S. M. Novel amyloid precursor protein mutation in an Iowa family with dementia and severe cerebral amyloid angiopathy. Ann. Neurol. (2001). doi:10.1002/ana.1009

30. Gould, D. B. et al. Role of COL4A1 in Small-Vessel Disease and Hemorrhagic Stroke. N. Engl. J. Med. (2006). doi:10.1056/NEJMoa053727

31. Lee, S. T. et al. A novel COL3A1 gene mutation in patient with aortic dissected aneurysm and cervical artery dissections. Heart Vessels (2008). doi:10.1007/s00380-007-1027-4

32. Shabbeer, J., Yasuda, M., Benson, S. D. & Desnick, R. J. Fabry disease: Identification of 50 novel α-galactosidase A mutations causing the classic phenotype and three-dimensional structural analysis of 29 missense mutations. Hum. Genomics (2006). doi:10.1186/1479-7364-2-5-297

33. Chinnery, P. F. & Hudson, G. Mitochondrial genetics. British Medical Bulletin (2013). doi:10.1093/bmb/ldt017

34. Steinberg, M. H. Predicting clinical severity in sickle cell anaemia. Br. J. Haematol. (2005). doi:10.1111/j.1365-2141.2005.05411.x

35. Milewicz, D. M. et al. De novo ACTA2 mutation causes a novel syndrome of multisystemic smooth muscle dysfunction. Am. J. Med. Genet. Part A (2010). doi:10.1002/ajmg.a.33657

36. Haberman, S., Capildeo, R. & Rose, F. C. Sex differences in the incidence of cerebrovascular disease. J. Epidemiol. Community Health 35, 45–50 (1981).

37. Russo, T., Felzani, G. & Marini, C. Stroke in the very old: A systematic review of studies on incidence, outcome, and resource use. Journal of Aging Research (2011). doi:10.4061/2011/108785

38. Kurth, T. et al. Body mass index and the risk of stroke in men. Arch. Intern. Med. (2002). doi:10.1001/archinte.162.22.2557

39. Aguet, F. et al. Genetic effects on gene expression across human tissues. Nature 550, 204–213 (2017).

40. Law, C. W., Chen, Y., Shi, W. & Smyth, G. K. Voom: Precision weights unlock linear model analysis tools for RNA-seq read counts. Genome Biol. 15, (2014).

41. Ritchie, M. E. et al. Limma powers differential expression analyses for RNA-sequencing and microarray studies. Nucleic Acids Res. 43, e47 (2015).

42. Fisher, R. A. Statistical methods for research workers. Biological monographs and manuals (1970). doi:10.1056/NEJMc061160

43. Baranova, O. et al. Neuron-specific inactivation of the hypoxia inducible factor 1α increases brain injury in a mouse model of transient focal cerebral ischemia. J. Neurosci. (2007). doi:10.1523/JNEUROSCI.0449-07.2007

44. Ashouri, A. et al. Pan-cancer transcriptomic analysis associates long non-coding RNAs with key mutational driver events. Nat. Commun. (2016). doi:10.1038/ncomms13197

45. Yamaguchi, Y. U. Brevican: A major proteoglycan in adult brain. Perspect. Dev. Neurobiol. (1996).

46. Ocak, U. et al. Targeting mast cell as a neuroprotective strategy. Brain Injury (2019). doi:10.1080/02699052.2018.1556807

47. Seifert, H. A. et al. Pro-Inflammatory Interferon Gamma Signaling is Directly Associated with Stroke Induced Neurodegeneration. J. Neuroimmune Pharmacol. (2014). doi:10.1007/s11481-014-9560-2

48. Gesuete, R., Kohama, S. G. & Stenzel-Poore, M. P. Toll-like receptors and ischemic brain injury. Journal of Neuropathology and Experimental Neurology (2014). doi:10.1097/NEN.0000000000000068

49. Smith, C. J. et al. Peak plasma interleukin-6 and other peripheral markers of inflammation in the first week of ischaemic stroke correlate with brain infarct volume, stroke severity and long-term outcome. BMC Neurol. (2004). doi:10.1186/1471-2377-4-2

50. Cresti, A. et al. Frequency and Significance of Right Atrial Appendage Thrombi in Patients with Persistent Atrial Fibrillation or Atrial Flutter. J. Am. Soc. Echocardiogr. (2014). doi:10.1016/j.echo.2014.08.008

51. Wang, T. et al. Effects of obesity related genetic variations on visceral and subcutaneous fat distribution in a Chinese population. Sci. Rep. (2016). doi:10.1038/srep20691

52. Chang, Y. C. et al. Common ALDH2 genetic variants predict development of hypertension in the SAPPHIRe prospective cohort: Gene-environmental interaction with alcohol consumption. BMC Cardiovasc. Disord. (2012). doi:10.1186/1471-2261-12-58

53. Kim, J. H. et al. Impact of visceral adipose tissue on clinical outcomes after acute ischemic stroke. Stroke (2019). doi:10.1161/STROKEAHA.118.023421

54. Danton, G. H. & Dietrich, W. D. Inflammatory mechanisms after ischemia and stroke. Journal of Neuropathology and Experimental Neurology (2003). doi:10.1093/jnen/62.2.127

55. Kawabori, M. & Yenari, M. Inflammatory Responses in Brain Ischemia. Curr. Med. Chem. (2015). doi:10.2174/0929867322666150209154036

56. Ock, J., Cho, H.-J., Hong, S., Kim, I. & Suk, K. Hypoxia as an Initiator of Neuroinflammation: Microglial Connections. Curr. Neuropharmacol. (2005). doi:10.2174/1570159053586681

57. Ferdinand, P. & Roffe, C. Hypoxia after stroke: a review of experimental and clinical evidence. Exp. Transl. Stroke Med. (2016). doi:10.1186/s13231-016-0023-0

58. Benita, Y. et al. An integrative genomics approach identifies Hypoxia Inducible Factor-1 (HIF-1)-target genes that form the core response to hypoxia. Nucleic Acids Res. (2009). doi:10.1093/nar/gkp425

59. Millwood, I. Y. et al. Conventional and genetic evidence on alcohol and vascular disease aetiology: a prospective study of 500 000 men and women in China. Lancet (2019). doi:10.1016/S0140-6736(18)31772-0

60. Rayasam, A. et al. Immune responses in stroke: How the immune system contributes to damage and healing after stroke and how this knowledge could be translated to better cures? Immunology (2018). doi:10.1111/imm.12918

61. Gelderblom, M. et al. Temporal and spatial dynamics of cerebral immune cell accumulation in stroke. Stroke (2009). doi:10.1161/STROKEAHA.108.534503

62. Krupinski, J., Kaluza, J., Kumar, P., Kumar, S. & Wang, J. M. Role of angiogenesis in patients with cerebral ischemic stroke. Stroke. (1994). doi:10.1161/01.STR.25.9.1794

63. Ho, S. Y., Anderson, R. H. & Sánchez-Quintana, D. Atrial structure and fibres: Morphologic bases of atrial conduction. Cardiovascular Research (2002). doi:10.1016/S0008-6363(02)00226-2

64. Ho, S. Y. & Sánchez-Quintana, D. The importance of atrial structure and fibers. Clinical Anatomy (2009). doi:10.1002/ca.20634

65. Al-Saady, N. M., Obel, O. A. & Camm, A. J. Left atrial appendage: Structure, function, and role in thromboembolism. Heart (1999). doi:10.1136/hrt.82.5.547

66. Subramaniam, B., Riley, M. F., Panzica, P. J. & Manning, W. J. Transesophageal echocardiographic assessment of right atrial appendage anatomy and function: comparison with the left atrial appendage and implications for local thrombus formation. J. Am. Soc. Echocardiogr. (2006). doi:10.1016/j.echo.2005.10.013

67. Paulson, J. N. et al. Tissue-aware RNA-Seq processing and normalization for heterogeneous and sparse data. BMC Bioinformatics 18, (2017).

68. Hicks, S. C. et al. Smooth quantile normalization. Biostatistics (2018). doi:10.1093/biostatistics/kxx028

69. Benjamini, Y. & Hochberg, Y. Controlling the false discovery rate: a practical and powerful approach to multiple testing. J. R. Stat. Soc. Ser. B (1995). doi:10.2307/2346101

70. Subramanian, A. et al. Gene set enrichment analysis: A knowledge-based approach for interpreting genome-wide expression profiles. Proc. Natl. Acad. Sci. (2005). doi:10.1073/pnas.0506580102

71. Köhler, S. et al. Expansion of the Human Phenotype Ontology (HPO) knowledge base and resources. Nucleic Acids Res. (2019). doi:10.1093/nar/gky1105

